# The anti-B7-H3 blocking antibody MJ18 does not recognize B7-H3 in murine tumor models

**DOI:** 10.1101/2023.11.15.567261

**Authors:** Talah Nammor, Jenna Frizzell, Roxane R. Lavoie, Fabrice Lucien

**Author notes:** **Corresponding Authors:** Fabrice Lucien, PhD, Mayo Clinic, Guggenheim 4-42B 200 1^St^ Street SW, Rochester, MN, 55901, USA. Contributed equally (Talah Nammor, Jenna Frizzell).

## Abstract

The immune checkpoint molecule B7-H3 is regarded as one of the most promising therapeutic targets for the treatment of human cancers. B7-H3 is highly expressed in many cancers and its expression has been associated to impaired antitumor immunity and poor patient prognosis. In immunocompetent mouse tumor models, genetic deletion of B7-H3 in tumor cells enhances antitumor immune response leading to tumor shrinkage. The underlying mechanisms of B7-H3 inhibitory function remain largely uncharacterized and the identity of potential cognate(s) receptor(s) of B7-H3 is still to be defined. To better understand B7-H3 function *in vivo*, several studies have employed MJ18, a monoclonal antibody reported to bind murine B7-H3 and blocks its immune-inhibitory function. In this brief research report, we show that 1) MJ18 does not bind B7-H3, 2) MJ18 binds the Fc receptor FcγRIIB on surface of murine splenocytes, and 3) MJ18 does not induce tumor regression in a mouse model responsive to B7-H3 knockout. Given the high profile of B7-H3 as therapeutic target for human cancers, our work emphasizes that murine B7-H3 studies using the MJ18 antibody should be interpreted with caution. Finally, we hope that our study will motivate the scientific community to establish much-needed validated research tools to study B7-H3 biology in mouse models.

## Introduction

The immune checkpoint molecule B7-H3 has become an attractive target for the development of novel cancer therapeutics^1^. While B7-H3 protein is rarely expressed in normal tissue, it is highly expressed in various cancers and elevated B7-H3 expression has been associated to poor prognosis^2, 3^. In tumors, B7-H3 has been mainly found on surface of tumor cells, tumor-associated endothelial cells and in some extent on surface of stroma cells^2, 4, 5^. For these reasons, B7-H3 is regarded as one of the most promising pan-cancer targets for antibody-drug conjugates^6^, bi- and tri-specific T/NK-cell engagers^7, 8^ and chimeric antigen receptor T cells^9^.

There is a growing literature on the role of B7-H3 in tumor immune evasion which provides a strong rationale to develop next-generation immune checkpoint inhibitors targeting B7-H3 in human cancers. B7-H3 expression has been linked to reduced tumor infiltration of NK and CD8 T cells and impaired cytotoxic activity^10–12^. In immunocompetent mouse models, genetic deletion of B7-H3 in tumor cells is associated with improved antitumor immunity and tumor shrinkage^11, 13, 14^. Despite this evidence, no B7-H3 immune checkpoint inhibitor has reached the clinic yet which can be explained by the fact that molecular mechanisms involved in the immunomodulatory role of B7-H3 remain largely uncharacterized and the identity of potential cognate(s) receptor(s) of B7-H3 is still to be defined. Furthermore, unlike other B7 molecules, mouse and human B7-H3 are structurally distinct in their extracellular domain due to exon duplication during evolution^15, 16^. While it remains to be determined, these distinct properties can confer to human B7-H3 unique functionalities compared to mouse B7-H3. In murine models, many studies have reported significant tumor regression in mice treated with MJ18, the only monoclonal antibody reported to bind murine B7-H3 and blocks its immune-inhibitory function (**Table 1**).

| Study | Year | Disease model | Dosage | Ref |
| --- | --- | --- | --- | --- |
| Fukushima A et al | 2007 | Allergic conjunctivitis | 300 µg/injection twice | 17 |
| Nagashima O et al | 2008 | Asthma | 300 µg/injection every 2-3 days | 18 |
| Yamato I et al | 2009 | Pancreatic cancer | 300 µg/injection every 2 days | 19 |
| Mao L et al | 2017 | Head and neck squamous cell carcinoma | 300 µg/injection every 2 days | 20 |
| Lee YH et al | 2017 | Lymphoma, Melanoma | 100 µg/injection every 2 days | 14 |
| Yonesaka K et al | 2018 | Pancreatic cancer | 15 mg/kg body weight per injection every 3 days | 13 |
| Cai D et al | 2020 | Ovarian cancer | 200 µg/injection every 2 days | 21 |
| Masemann D et al | 2021 | Lung cancer | 250 µg/injection every 3 days | 22 |
| Zhou Z et al | 2022 | Hepatocellular carcinoma | Not available | 23 |
| Liu T et al | 2022 | Pulmonary fibrosis | 300 µg/injection every 2 days | 24 |
| Shi W et al | 2023 | Prostate cancer | 300 µg/injection every 3 days | 11 |

The MJ18 antibody has been first described by Nagashima et. al^18^. MJ18 was generated by immunization of Sprague Dawley rats with a fusion protein containing the extracellular domain of murine B7-H3 fused to a Fc portion of mouse IgG2a. Binding of MJ18 to mouse B7-H3 was tested by flow cytometry with B7-H3-overexpressing P815 mastocytoma cells and L5178Y lymphoma cells. No cross-reactivity with other members of the B7 family was reported.

Our group recently identified B7-H3 as an important mediator of immune evasion in rhabdomyosarcoma, the most common soft-tissue sarcoma in children^10^. To dissect the biology of B7-H3 in rhabdomyosarcoma, we sought to determine the antitumor activity of MJ18 in a syngeneic mouse model of rhabdomyosarcoma (M3-9-M). Unfortunately, our data indicate that MJ18 does not recognize B7-H3 on surface of murine tumor cell lines but Fc receptors on surface of murine splenocytes. These findings cast doubt on the value and reliability of published studies using the MJ18 antibody clone. In this expression of concern, we aim to alert the scientific community of our controversial data and encourage validation studies for the target(s) of MJ18 and its functional impact *in vivo*. Finally, we hope our work will raise awareness of the lack of reliable tools to study murine B7-H3 biology and translate discoveries into clinical applications.

## Results

### MJ18 does not bind B7-H3 but recognizes a ligand on surface of murine splenocytes

To validate the specific binding of MJ18 to murine B7-H3, we first determined the protein expression of B7-H3 in various tumor cell lines by western-blot. We used the EPNCIR122 clone, a rabbit monoclonal antibody that recognizes both mouse and human B7-H3^10, 25^. Murine tumor cell lines M3-9-M (rhabdomyosarcoma) and RM-1 (prostate cancer) showed B7-H3 expression (molecular weight ~50-60 kDa) which was lost following gene deletion with CRISPR/Cas9 technology (**Figure 1A**). M3-9-M transfected either with murine or human B7-H3 revealed bands at expected molecular weights, in other words ~50-60 kDa for murine B7-H3 and ~98 kDa for its human counterpart. This corroborates with the structural differences between both species^16^. Expression of human B7-H3 was detectable in human tumor cell lines RH30 (rhabdomyosarcoma) and PC3 (prostate cancer). No band was detected in tumor cells knockout for B7-H3 which confirms the specificity of EPNCIR122 for B7-H3. B7-H3 expressing and knockout murine tumor cells (M3-9-M, RM-1, TRAMP-C2,) were analyzed for the binding of EPNCIR122 and MJ18 by flow cytometry. All wild-type cell lines tested showed positive signal with EPNCIR122 which was lost in B7-H3 knockout cells (**Figure 1B**). In contrast, no positive binding of MJ18 was detected for the three cell lines. To our knowledge, it represents the first attempt to validate the binding of MJ18 on B7-H3 by flow cytometry.

**Figure 1.**
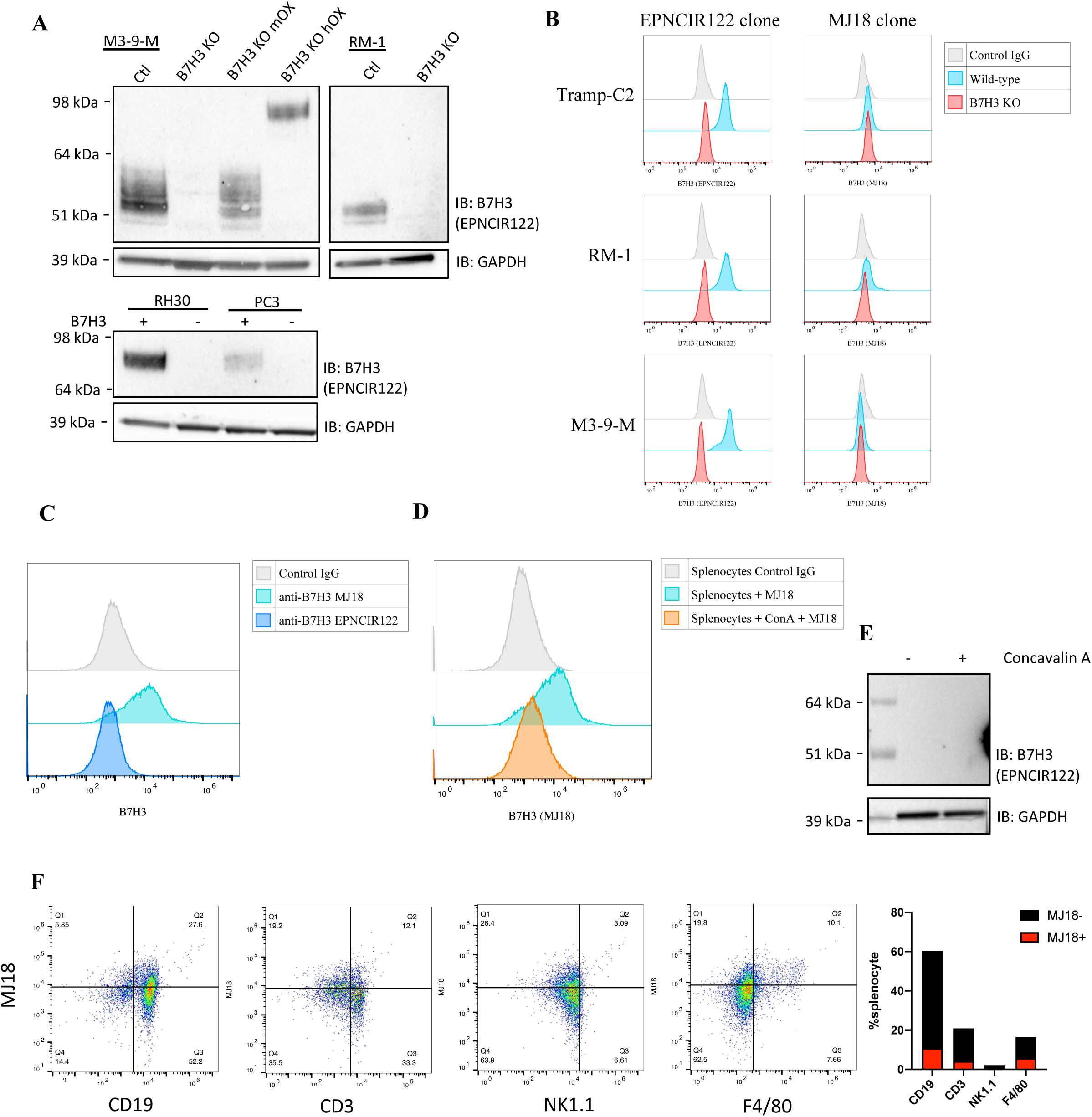
MJ18 does not bind B7-H3 but recognizes a ligand on surface of murine splenocytes. **A)** Western blot showing B7-H3 expression in mouse and human tumor cell lines. Predicted molecular weight for mouse and human B7-H3 is ~50-90 kDa and ~90-100 kDa. Anti-B7-H3 EPNCIR122 clone was used to detect both mouse and human B7-H3. GAPDH was used as loading control. **B)** Surface expression of B7-H3 in wild-type and B7-H3 knockout tumor cell lines (Tramp-C2: prostate; RM-1: prostate; M3-9-M: rhabdomyosarcoma) measured by flow cytometry with EPNCIR122 and MJ18 clones. **C)** Binding of EPNCIR122 and MJ18 antibodies on surface of murine splenocytes from C57/BL6 wild-type mice. **D)** Binding of MJ18 antibodies on surface of murine splenocytes treated with Concavalin A. **E)** Western blot showing the lack of B7-H3 expression in murine splenocytes using the EPNCIR122 antibody clone. **F)** Scatterplots showing the immune cell origin of MJ18 positivity using CD19 (B cells), CD3 (T cells), NK1.1 (NK cells) and F4/80 (macrophages) markers. Bar graph summarizes the immune cell origin of MJ18 binding.

Murine B7-H3 was initially discovered as an inducible surface molecule on surface of dendritic cells, monocytes and T cells^26^ and B7-H3 binding by MJ18 was initially tested by flow cytometry on mastocytoma and lymphoma cells^18^. Therefore, we sought to assess the binding of MJ18 on murine splenocytes. Flow cytometry revealed a positive signal with 30% of splenocytes positive for MJ18 (**Figure 1C**). Interestingly, MJ18 binding was lost in murine splenocytes stimulated overnight with concavalin A, a mitogen that induces proliferation of T cells (**Figure 1D**). Loss of the MJ18 signal suggests that T-cells are not the main cellular source of MJ18 ligand or that concavalin A reduces surface expression of the MJ18 ligand. The EPNCIR122 did not show any positive signal in both untreated and concavalin A-treated splenocytes by flow cytometry and western-blot (**Figure 1C-1E**). Immunophenotyping identified B cells (CD19+) macrophages (F4/80^+^) and T cells (CD3^+^) as MJ18-positive cells (**Figure 1F**). These data demonstrate that MJ18 bind a protein on surface of murine splenocytes, but it is not identified to be B7-H3.

### Immunoprecipitation-Mass Spectrometry (IP-MS) rejects MJ18 as an antibody against B7-H3

To reveal the identity of MJ18 ligand, we performed protein immunoprecipitation followed by mass spectrometry (IP-MS). To that end, M3-9-M tumor cells and splenocytes were labeled with biotinylated EPNCIR122 or MJ18 antibodies prior to streptavidin bead-based immunoprecipitation (**Figure 2A**). By western-blot, B7-H3 was only detected in M3-9-M tumor cells labeled with ENPCIR122 only. The identity of tumor- and splenocyte-derived proteins isolated by IP with MJ18 and EPNCIR122 was determined by mass spectrometry. IP-MS can reveal multiple false-positive hits which defined unspecific binding of abundant protein contaminants to the IP reagents^27^. To minimize the false-discovery rate, we annotated proteins detected to the CRAPome, a repository of background contaminants commonly found in IP-MS experiments (**Figure 2C**). In our IP-MS experiments, ~80-85% of proteins detected were background contaminants resulting in a final protein list of 71 proteins. Starting with tumor cells, we determined differences in protein repertoire between methods of pulldown, in other words EPN, MJ18, No Ab (**Figure 2D**). Ten proteins were commonly shared by TC_EPN and TC_MJ18 samples and it included B7-H3. In the TC_EPN sample, B7-H3 (CD276) was the most abundant protein (**Figure 2E**). The second most abundant protein NCCRP1 had expression level 50.2 times lower than B7-H3. Despite B7-H3 was found in the TC_MJ18 sample, its protein expression was 44.7 lower than in the TC_EPN sample. NCCRP1 was also the second most abundant protein in the TC_MJ18 sample and its expression was only 1.12 times lower than B7-H3. Three and two proteins were found uniquely in TC_EPN and TC_MJ18 respectively (**Figure 2D**). Their level of expression remains in the range of unspecific binders found in both samples (**Figure 2F**). IP-MS demonstrates that B7-H3 stands out as the primary target of the EPNCIR122 antibody clone in tumor cells. It also confirms that B7-H3 is not the target of the MJ18 antibody clone.

**Figure 2.**
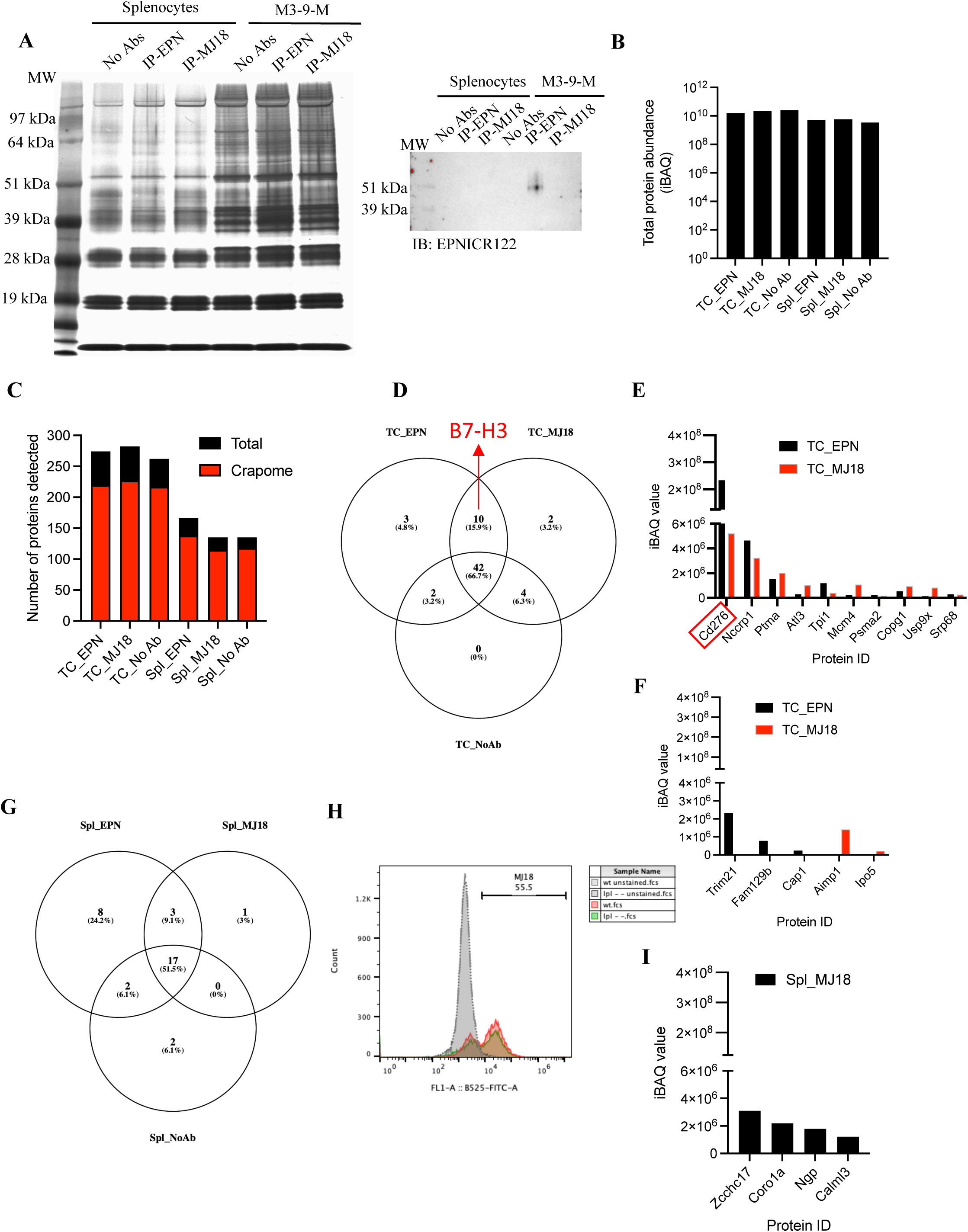
Immunoprecipitation-Mass Spectrometry (IP-MS) rejects MJ18 as an antibody against B7-H3. **A)** Silver staining of polyacrylamide gel for detection of proteins pulled-down by immunoprecipitation with EPNCIR122 or MJ18 clones in both M3-9-M tumor cells and murine splenocytes. Western blot shows detection of mouse B7-H3 following IP with EPNCIR122 in M3-9-M tumor cells. **B)** Total abundance values (iBAQ) following mass spectrometry analysis of IP with EPNCIR122 or MJ18 clones in both M3-9-M tumor cells and murine splenocytes. **C)** Total number of proteins detected and number of proteins associated to the Crapome **D)** Venn diagram showing differences in protein repertoire obtained from IP with EPNCIR122 and MJ18 in tumor cells. No antibody IP was used as negative control. **E)** Abundance of proteins identified in IP with both EPNCIR122 and MJ18. Red rectangle highlights B7-H3 (*cd276*) **F)** Abundance of proteins uniquely identified in IP with EPNCIR122 (dark) or MJ18 (red). **G)** Venn diagram showing differences in protein repertoire obtained from IP with EPNCIR122 and MJ18 in murine splenocytes. No antibody IP was used as negative control. **H)** Histogram showing MJ18 binding in splenocytes from wild-type and LPL^-/-^ C57BL/6. **I)** Abundance of proteins identified in IP with MJ18 in murine splenocytes but not in tumor cells.

In splenocytes, 21 proteins were identified with MJ18 pulldown. Seventeen of them were shared with EPN and NoAb samples and three with EPN alone (**Figure 2G**). One protein, Lymphocyte Cytosolic Protein 1 (LCP1), was found only in the MJ18 sample. LCP1 (L-Plastin, Plastin 2) is an actin-bundling protein important in the formation of the cytolytic synapse and cytotoxic function of T cells^28^. LCP1 is also found on surface of B cells and participates to chemotaxis and motility^29^. Based on our immunophenotyping data showing B cells as the major MJ18-positive population, LCP1 appears as a potential ligand for MJ18. To validate this, we compared the binding of MJ18 on surface of splenocytes isolated from LCP1^-/-^ mice^30^. We did not find any difference in the binding of MJ18 between wild-type and LCP1^-/-^ mice-derived splenocytes (**Figure 2H**). Among the others 20 proteins found in Spl_MJ18 samples, only 4 (ZCCHC17, CORO1A, NGP, CALML3) were not detected in TC_MJ18 (**Figure 2I**). However, their expression levels were in the range of unspecific binders found in tumor cells.

Since we did not find any specific binder of MJ18, we sought to determine whether MJ18 can be bound by Fc receptors expressed on surface of murine splenocytes. To that end, we labeled murine splenocytes with MJ18 with or without pretreatment with a Fc blocking agent (anti-CD16/32). Unexpectedly, we observed a strong reduction of the fluorescence signal associated to MJ18 intensity with ~60% of MJ18-positive cells and ~10% following Fc block (**Figure 3A**). IP-MS confirmed MJ18 as a non-B7-H3 antibody and flow cytometry identified Fc receptors as the specific binder of MJ18.

**Figure 3.**
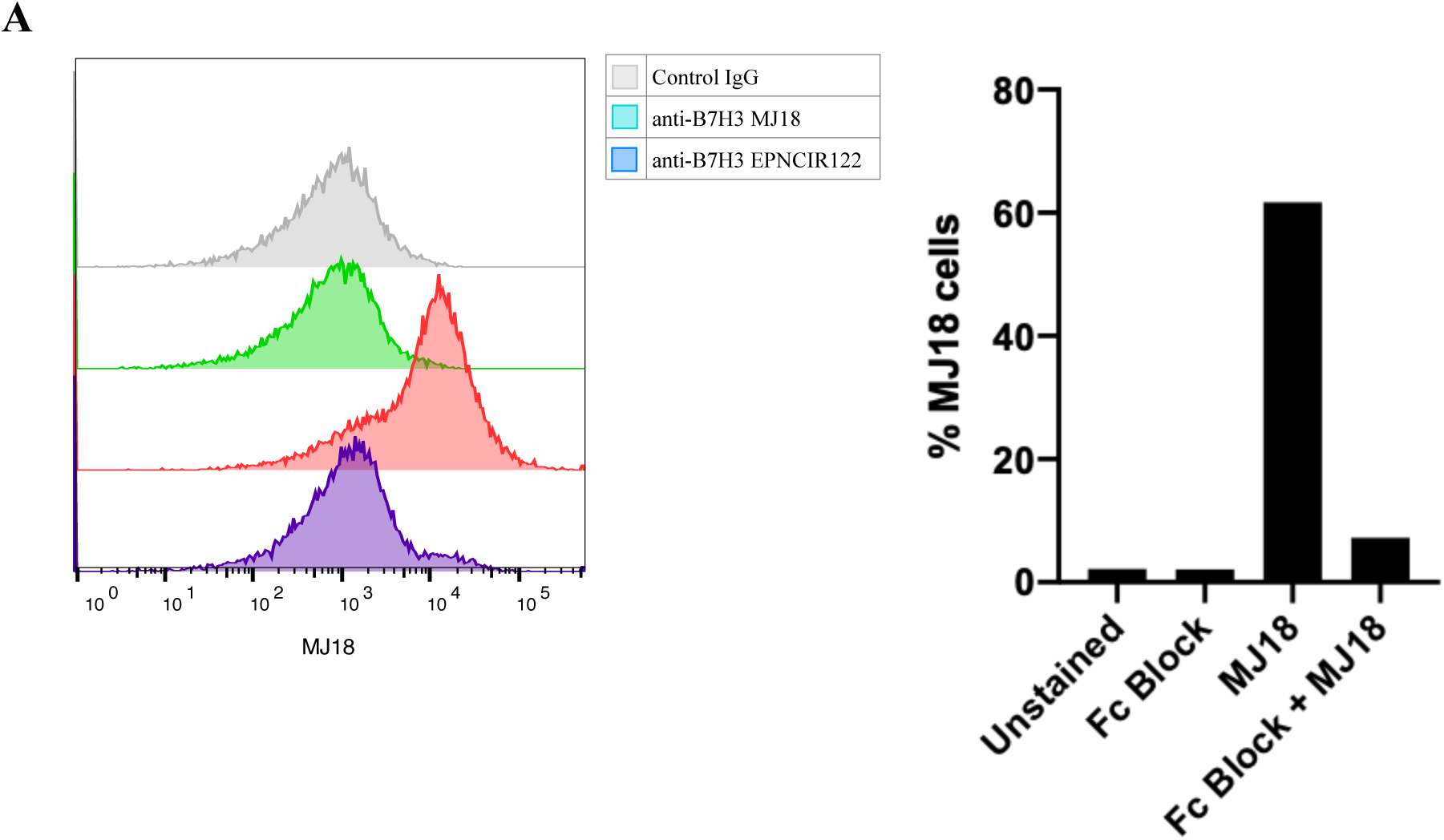
MJ18 binds Fc receptors on surface of murine splenocytes. **A)** Density plots showing MJ18 binding on murine splenocytes treated or not with Fc block. Bar graph indicates the impact of FC block on MJ18 binding in murine splenocytes

### MJ18 does not induce tumor regression in a murine B7-H3 expressing tumor model

Several studies have reported an antitumor activity of MJ18 in mouse syngeneic tumor models (**Table 1**). Although B7-H3 is not the target of MJ18, tumor regression and increased infiltration of T cells in MJ18-treated mice may shed the light on an uncharacterized yet mechanism of antitumor immune response. To validate prior findings, we tested the antitumor activity of MJ18 in an orthotopic mouse model of rhabdomyosarcoma (M3-9-M)^31^. B7-H3 knockout tumors were used as positive control. M3-9-M is a poorly immunogenic fast-growing tumor model. Following subcutaneous injection of 0.5×10^6^ cells, mice bearing wild-type M3-9-M tumors reached humane endpoint by Day 13 (**Figure 4A**). In contrast, B7-H3 knockout tumors showed tumor growth delay compared to wild-type tumors (**Figure 4B**) which validates M3-9-M as a suitable tumor model responsive to B7-H3 inhibition. To assess the antitumor activity of MJ18, we inoculated 0.3×10^6^ cells which led to better tumor control in B7-H3^+^ tumor-bearing mice. Treatment with MJ18 antibody using the most reported regimen (300 µg per injection every other day) did not show any difference in tumor growth with vehicle-treated mice (**Figure 4C**). While B7-H3 knockout improves tumor control in vivo, it was not recapitulated with the MJ18 antibody.

**Figure 4.**
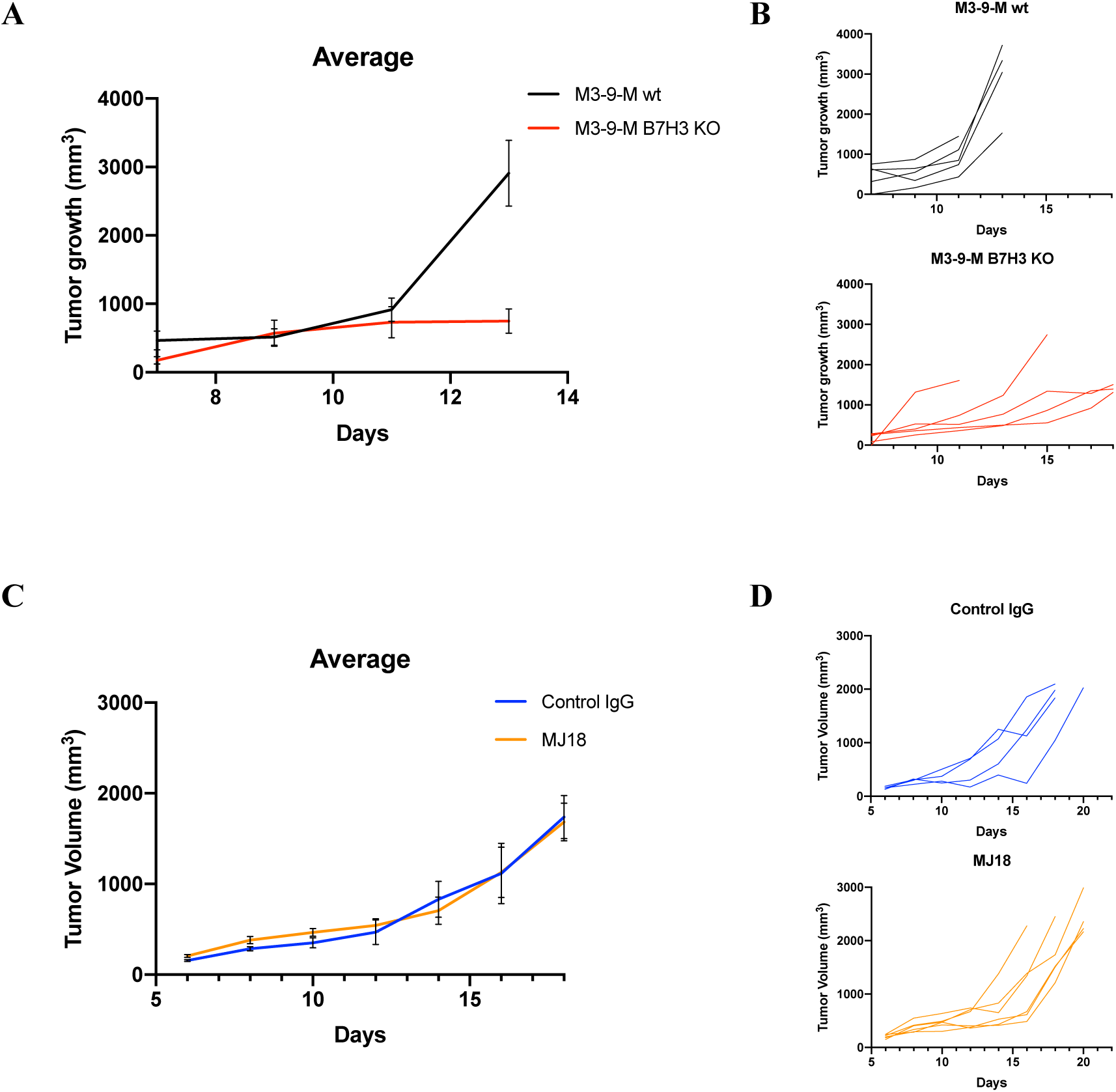
MJ18 does not induce tumor regression in a murine B7-H3 expressing tumor model. **A-B)** Grouped (**A**) and individual (**B**) tumor growth curves for wild-type and B7-H3 knockout M3-9-M-bearing mice. Statistical significance for differences in tumor growth at Day 13 was determined by Mann-Whitney test, p=0.029. **C-D)** Grouped (**C**) and individual (**D**) tumor growth curves for wild-type M3-9-M-bearing mice treated with MJ18 or IgG control.

## Discussion

In this study, we shed the light on an erroneous utilization of the MJ18 clone as a blocking antibody targeting murine B7-H3. Using flow cytometry analysis of a variety of murine genetically modified tumor cells and splenocytes, we demonstrated that MJ18 does not recognize B7-H3 on surface of tumor cells but bind an unidentified ligand on surface of murine splenocytes with preference for B cells. Surprisingly, we did not find any reported data validating MJ18 binding on surface of B7-H3-expressing cells.

We also validated the clone EPNCIR122 as a specific antibody for both murine and human B7-H3. The EPNCIR122 helped us claimed that murine splenocytes from healthy C57BL/6 mice do not express B7-H3 on their surface. We performed IP-MS to further validate our flow cytometry data and identify putative targets of MJ18 in murine splenocytes. IP-MS confirmed EPNCIR122 as a B7-H3-specific antibody and MJ18 as an “orphan” antibody. We identified several surface proteins that could be the target of MJ18 but their low expression is similar to unspecific binders and does not clearly show an unequivocal target. In fact, we found Fc receptors as specific binders of MJ18 on surface of murine splenocytes. Five Fc receptors Fc_γ_RI, Fc_γ_RIIB, Fc_γ_RIII, Fc_γ_RIV, FcRn are expressed across immune cells in mice^32^. A recent study revealed that rat IgG1 binds exclusively with Fc_γ_RIIB and Fc_γ_RIII^33^. Mouse Fc_γ_RIIB is the only inhibitory Fc receptor expressed by murine splenic B cells, the major immune cell subset bound by MJ18^32^. Fc_γ_RIIB can also be found on surface of T cells, another type of immune cells positive for MJ18^34^. While our study goal is to demonstrate that MJ18 does not bind B7-H3, validation Fc_γ_RIIB as MJ18 target using Fc_γ_RIIB^-/-^ mice is warranted.

Using a syngeneic mouse tumor model of rhabdomyosarcoma, we confirmed that B7-H3 knockout significantly delays tumor growth, in accordance with the reported role of B7-H3 as an inhibitory immune checkpoint molecule. In contrast, we did not find any antitumor activity of MJ18 at a similar dose previously reported. The paradoxical response observed with MJ18 cast doubt on the validity and reproducibility of the reported antitumor activity of the MJ18 antibody in mouse tumor models. By providing a transparent validation of MJ18 as an antibody not specific to B7-H3 with no antitumor activity, we emphasize the critical need to better validate research tools and report both positive and negative findings before it leads to data misinterpretation and irreproducibility. Such rigorous and iterative process is instrumental for the translation of bench discoveries to the bedside. Unfortunately, the lack of specificity of antibodies used in research is not new and it has already been under the spotlight^35, 36^. This alarming situation was used to raise awareness and develop government funded programs with the mission of creating public resources of validated antibodies for multiple downstream applications^37^. In parallel, guidelines have been established for research groups to validate antibody specificity and ensure their reproducibility^38^.

To conclude, we conducted a rigorous evaluation of the MJ18 antibody and found that B7-H3 is not the target of MJ18. We also did not observe any antitumor activity of MJ18 *in vivo*. Novel and validated B7-H3 are critically needed to better understand the mechanistic basis of B7-H3 immune function using murine models.

## Materials and Methods

### Human and murine cells

Human rhabdomyosarcoma (RH30) and prostate cancer (PC3) cell lines were obtained from ATCC (Manassas, VA, USA) and maintained in a humidified incubator at 5% CO_2_. RH3O and PC3 cells were cultured in RPMI 1640 (Corning, New York, NY, USA) with 10% fetal bovine serum (Thermo Fisher Scientific, Waltham, MA) and 1% penicillin and streptomycin (Thermo Fisher Scientific, Waltham, MA). Mouse rhabdomyosarcoma (M3-9-M) was obtained from Dr. Crystal Mackall (Stanford University, Stanford, CA)^31^ and cultured in RPMI 1640 (Corning, New York, NY, USA) with 10% fetal bovine serum (Thermo Fisher Scientific, Waltham, MA), 1% HEPES buffer (Thermo Fisher Scientific, Waltham, MA), 1% non-essential amino acids (Thermo Fisher Scientific, Waltham, MA), 1% sodium pyruvate (Thermo Fisher Scientific, Waltham, MA) and 1% penicillin and streptomycin (Thermo Fisher Scientific, Waltham, MA). Murine prostate cancer (RM-1 and TrampC2) cell lines were obtained from ATCC (Manassas, VA, USA) and maintained in a humidified incubator at 5% CO_2_. RM-1 and TrampC2 cells were cultured in DMEM (Corning, New York, NY, USA) with 10% fetal bovine serum (Thermo Fisher Scientific, Waltham, MA) and 1% penicillin and streptomycin (Thermo Fisher Scientific, Waltham, MA).

Murine splenocytes were isolated from spleens of C57BL/6 wild-type mice. Spleens were dissociated and cell suspensions were treated with ACK buffer to remove red blood cells. Splenocytes were stimulated overnight with Concanavalin A (5 µg/mL, Sigma-Aldrich, Saint-Louis, MO) in RPMI 1640 (Corning, New York, NY, USA) supplemented with 10% fetal bovine serum (Thermo Fisher Scientific, Waltham, MA) and 1% penicillin and streptomycin (Thermo Fisher Scientific, Waltham, MA) and maintained in a humidified incubator at 5% CO_2_. Cryopreserved murine splenocytes from L-Plastin knockout mice (LCP1^-/-^) mice were kindly provided by Dr. Sharon Celeste Morley (Washington University School of Medicine, Saint-Louis, MO)^30^. LCP1^-/-^ murine splenocytes were thawed at 37C and prepared as described above.

### B7-H3 CRISPR/Cas9 Knockout

B7-H3 (*cd276*) was knocked-out in tumor cell lines using CRISPR/Cas9 KO plasmids from Santa Cruz Biotechnology (human: sc-402032, mouse: sc-430440). Human and tumor cell lines were transfected with CRISPR/Cas9 KO plasmids using lipofectamine 3000 and manufacturer’s instructions. Twenty-four hours after transfection, GFP-positive cells were sorted by FACS. A second round of sorting were conducted using B7-H3 antibody staining (EPNCIR122 clone, Abcam, ab134161) to obtain a polyclonal population of B7-H3 knockout cells. B7-H3 knockout was validated by flow cytometry and western blot.

### Flow cytometry

Tumor cell lines and murine splenocytes were stained for flow cytometry assays in FACS buffer (1X PBS, 2 mM EDTA and 2c% FBS) with the following antibody panel: CD3-BV510 (BioLegend, clone 17A2, cat. 100233), NK1.1-PE-Cy7 (BioLegend, clone PK136, cat. 108713), CD19-FITC (BioLegend, clone 6D5, cat. 115505), F4/80-PE (BioLegend, clone BM8, cat. 123109), B7-H3-AF647 (Abcam, clone EPNCIR122, ab134161), B7-H3-AF647 (BioXcell, clone MJ18, catalog BE0124). When indicated, cells were treated with Fc block prior to antibody staining (TruStain FcX™ PLUS (anti-mouse CD16/32) Antibody, BioLegend, San Diego, CA). Data acquisition was performed on a CytoFLEX LX (Beckman Coulter) and analyzed with FlowJo 10.9 (BD Biosciences).

### Western blot

Whole-cell extracts were lysed by radioimmunoprecipitation assay buffer (Thermo Scientific, 89901) containing protease inhibitors (Thermo Scientific, 1860932) and protein concentration was determined using BCA assay (Thermo Scientific, 23228). Immunoblotting was performed using a 4-12% gradient precast gel (Invitrogen, NP0321BOX) and nitrocellulose membrane using a semi-dry blotting system (Invitrogen, IB23002). The membrane was blocked in TBS containing 3% BSA for 1h at RT and incubated with rabbit anti-B7H3 (1:1000, clone EPNCIR122, Abcam, ab134161) and rabbit anti-GAPDH (1:1000, Cell Signaling, 5174S) O/N at 4^0^C. After washing the membrane with TBS/0.1%Tween-20, membrane was incubated with HRP-conjugated secondary antibody (1:10,000, Invitrogen, 65-6120). Protein detection was performed using chemiluminescent substrate (Thermo Scientific, 34580) and a iBright Imaging System Biorad (Thermo Scientific).

### Immunoprecipitation and mass spectrometry (IP-MS)

Cultured tumor cells were trypsinized, washed and resuspended in ice-cold PBS. Tumor cells and murine splenocytes were incubated with biotinylated anti-B7-H3 (EPNCIR122 or MJ18) for 30 minutes on ice. The cells were washed with 50 mM of glycine to quench the unbound biotin. The cells were lysed in NP-40 lysis buffer, and the biotinylated cell surface proteins were affinity-purified on streptavidin magnetic beads (Thermo Scientific, cat#88816). After stripping off the nonspecifically bound proteins by several rounds of washing with the lysis buffer, the labeled proteins were reduced with 10-mM TCEP (Thermo Scientific, cat#77720) for 30 min at 50 _◦_C and followed by alkylation with iodoacetamide (Thermo Scientific, 90034) for 30 min in the dark at room temperature. A fraction of the protein lysate was run in a SDS-PAGE electrophoresis gel and silver stained (Thermo, 24612). The remaining protein lysate was submitted for mass spectrometry (Proteomics Core). Samples were digested with Micro S-Traps (Protifi). The peptide extracts (1 ug) were analyzed by nano-ESI-LC/MS/MS with an Exploris mass spectrometer coupled to a dionex nano-LC system (Thermo Scientific; Waltham, MA). The LC system used a gradient with solvent A (2% ACN, 0.2% formic acid, in water) and solvent B (80% ACN, 10% IPA, 0.2% formic acid, in water) as follows: −4-5 min, 5% B; 5-125 min 5-45 % gradient; 125-128 min 45-95% gradient; 128-132 min 95% B; 132-134 min 95-5% B gradient; 134-137 min 5% B with a flow rate of 300nL/min. The mass spectrometer had a resolution of 60,000 at 200 m/z and used data dependent acquisition, with a full MS1 scan ranging from 340-1600 m/z. Dynamic exclusion was set to 25 seconds. Cycle time was 3 seconds. All MS/MS spectra were analyzed using MaxQuant (Max Planck Institute of Biochemistry, Version 1.6.17.0). Each was set up to search the current UniProt database, assuming trypsin digestion with up to two miscleavages with a fragment ion tolerance of 20 PPM and parent ion tolerance of 4.5 ppm (UniProt is provided in the public domain by the Swiss Institute of Bioinformatics, Geneva, Switzerland, www.uniprot.org). Oxidation of methionine was set as a variable modification, and carbamidomethylation of cysteine (iodoacetamide derivative) was set as a fixed modification.

### In vivo tumor models

C57BL/6 mouse were purchased from Jackson Laboratory. All animals were housed under standard conditions (21 – 22 °C; 55% humidity) in individually ventilated cages, with a 12-h light/dark cycle and ad libitum access to food and water. Mice aged between 8 to 14 weeks were used in the studies. All experimental procedures were approved by the Mayo Clinic’s Institutional Animal Care and Use Committee (IACUC). M3-9-M cells were harvested by trypsin digestion, a single cell suspension was prepared, mice were shaved on the left hindlimb area and tumor cells (5×10^5^ or 3×10^5^ cells) were injected into the gastrocnemius muscle in 0.1 ml PBS using a sterile 27.5 G needle.

### Statistical Analysis

Student *t*-test (parametric), Mann-Whitney test (non-parametric) was employed to compare two groups. One-way ANOVA (parametric) and Kruskal-Wallis (non-parametric) tests were used to compare three or more groups. The results were considered significant for p values <0.05. p values were either specified in the figure or denoted as asterisk: * p<0.05, ** p<0.01, *** p<0.001, **** p<0.0001. All data were analyzed and plotted in GraphPad Prism 9.5.0.

## Acknowledgments

We acknowledge the assistance of the Mayo Clinic Proteomics Core, which is a shared resource of the Mayo Clinic Cancer Center (NCI P30 CA15083).

## Author Contributions

RRL and FL designed experiments. TN, JF, RRL performed experiments. TN, JF, RRL and FL wrote the manuscript. FL conceptualized and supervised the study.

## References

1. Liu C, Zhang G, Xiang K, Kim Y, Lavoie RR, Lucien F, Wen T. Targeting the immune checkpoint B7-H3 for next-generation cancer immunotherapy. Cancer Immunol Immunother. 2022;71(7):1549–67. Epub 20211105. doi: 10.1007/s00262-021-03097-x. PubMed PMID: 34739560.

2. Crispen PL, Sheinin Y, Roth TJ, Lohse CM, Kuntz SM, Frigola X, Thompson RH, Boorjian SA, Dong H, Leibovich BC, Blute ML, Kwon ED. Tumor cell and tumor vasculature expression of B7-H3 predict survival in clear cell renal cell carcinoma. Clin Cancer Res. 2008;14(16):5150–7. Epub 20080811. doi: 10.1158/1078-0432.CCR-08-0536. PubMed PMID: 18694993; PMCID: PMC2789387.

3. Roth TJ, Sheinin Y, Lohse CM, Kuntz SM, Frigola X, Inman BA, Krambeck AE, McKenney ME, Karnes RJ, Blute ML, Cheville JC, Sebo TJ, Kwon ED. B7-H3 ligand expression by prostate cancer: a novel marker of prognosis and potential target for therapy. Cancer Res. 2007;67(16):7893–900. Epub 2007/08/10. doi: 10.1158/0008-5472.CAN-07-1068. PubMed PMID: 17686830.

4. Seaman S, Zhu Z, Saha S, Zhang XM, Yang MY, Hilton MB, Morris K, Szot C, Morris H, Swing DA, Tessarollo L, Smith SW, Degrado S, Borkin D, Jain N, Scheiermann J, Feng Y, Wang Y, Li J, Welsch D, DeCrescenzo G, Chaudhary A, Zudaire E, Klarmann KD, Keller JR, Dimitrov DS, St Croix B. Eradication of Tumors through Simultaneous Ablation of CD276/B7-H3-Positive Tumor Cells and Tumor Vasculature. Cancer Cell. 2017;31(4):501–15 e8. doi: 10.1016/j.ccell.2017.03.005. PubMed PMID: 28399408; PMCID: PMC5458750.

5. MacGregor HL, Sayad A, Elia A, Wang BX, Katz SR, Shaw PA, Clarke BA, Crome SQ, Robert-Tissot C, Bernardini MQ, Nguyen LT, Ohashi PS. High expression of B7-H3 on stromal cells defines tumor and stromal compartments in epithelial ovarian cancer and is associated with limited immune activation. J Immunother Cancer. 2019;7(1):357. Epub 20191231. doi: 10.1186/s40425-019-0816-5. PubMed PMID: 31892360; PMCID: PMC6937725.

6. Scribner JA, Brown JG, Son T, Chiechi M, Li P, Sharma S, Li H, De Costa A, Li Y, Chen Y, Easton A, Yee-Toy NC, Chen FZ, Gorlatov S, Barat B, Huang L, Wolff CR, Hooley J, Hotaling TE, Gaynutdinov T, Ciccarone V, Tamura J, Koenig S, Moore PA, Bonvini E, Loo D. Preclinical Development of MGC018, a Duocarmycin-based Antibody-drug Conjugate Targeting B7-H3 for Solid Cancer. Mol Cancer Ther. 2020;19(11):2235–44. Epub 20200923. doi: 10.1158/1535-7163.MCT-20-0116. PubMed PMID: 32967924.

7. Vallera DA, Ferrone S, Kodal B, Hinderlie P, Bendzick L, Ettestad B, Hallstrom C, Zorko NA, Rao A, Fujioka N, Ryan CJ, Geller MA, Miller JS, Felices M. NK-Cell-Mediated Targeting of Various Solid Tumors Using a B7-H3 Tri-Specific Killer Engager In Vitro and In Vivo. Cancers (Basel). 2020;12(9). Epub 20200918. doi: 10.3390/cancers12092659. PubMed PMID: 32961861; PMCID: PMC7564091.

8. You G, Lee Y, Kang YW, Park HW, Park K, Kim H, Kim YM, Kim S, Kim JH, Moon D, Chung H, Son W, Jung UJ, Park E, Lee S, Son YG, Eom J, Won J, Park Y, Jung J, Lee SW. B7-H3x4-1BB bispecific antibody augments antitumor immunity by enhancing terminally differentiated CD8(+) tumor-infiltrating lymphocytes. Sci Adv. 2021;7(3). Epub 20210115. doi: 10.1126/sciadv.aax3160. PubMed PMID: 33523913; PMCID: PMC7810375.

9. Theruvath J, Sotillo E, Mount CW, Graef CM, Delaidelli A, Heitzeneder S, Labanieh L, Dhingra S, Leruste A, Majzner RG, Xu P, Mueller S, Yecies DW, Finetti MA, Williamson D, Johann PD, Kool M, Pfister S, Hasselblatt M, Fruhwald MC, Delattre O, Surdez D, Bourdeaut F, Puget S, Zaidi S, Mitra SS, Cheshier S, Sorensen PH, Monje M, Mackall CL. Locoregionally administered B7-H3-targeted CAR T cells for treatment of atypical teratoid/rhabdoid tumors. Nat Med. 2020;26(5):712–9. Epub 20200427. doi: 10.1038/s41591-020-0821-8. PubMed PMID: 32341579; PMCID: PMC7992505.

10. Lavoie RR, Gargollo PC, Ahmed ME, Kim Y, Baer E, Phelps DA, Charlesworth CM, Madden BJ, Wang L, Houghton PJ, Cheville J, Dong H, Granberg CF, Lucien F. Surfaceome Profiling of Rhabdomyosarcoma Reveals B7-H3 as a Mediator of Immune Evasion. Cancers (Basel). 2021;13(18). Epub 2021/09/29. doi: 10.3390/cancers13184528. PubMed PMID: 34572755; PMCID: PMC8466404.

11. Shi W, Wang Y, Zhao Y, Kim JJ, Li H, Meng C, Chen F, Zhang J, Mak DH, Van V, Leo J, St Croix B, Aparicio A, Zhao D. Immune checkpoint B7-H3 is a therapeutic vulnerability in prostate cancer harboring PTEN and TP53 deficiencies. Sci Transl Med. 2023;15(695):eadf6724. Epub 20230510. doi: 10.1126/scitranslmed.adf6724. PubMed PMID: 37163614.

12. Miyamoto T, Murakami R, Hamanishi J, Tanigaki K, Hosoe Y, Mise N, Takamatsu S, Mise Y, Ukita M, Taki M, Yamanoi K, Horikawa N, Abiko K, Yamaguchi K, Baba T, Matsumura N, Mandai M. B7-H3 Suppresses Antitumor Immunity via the CCL2-CCR2-M2 Macrophage Axis and Contributes to Ovarian Cancer Progression. Cancer Immunol Res. 2022;10(1):56–69. Epub 20211119. doi: 10.1158/2326-6066.CIR-21-0407. PubMed PMID: 34799346; PMCID: PMC9414298.

13. Yonesaka K, Haratani K, Takamura S, Sakai H, Kato R, Takegawa N, Takahama T, Tanaka K, Hayashi H, Takeda M, Kato S, Maenishi O, Sakai K, Chiba Y, Okabe T, Kudo K, Hasegawa Y, Kaneda H, Yamato M, Hirotani K, Miyazawa M, Nishio K, Nakagawa K. B7-H3 Negatively Modulates CTL-Mediated Cancer Immunity. Clin Cancer Res. 2018;24(11):2653–64. Epub 20180312. doi: 10.1158/1078-0432.CCR-17-2852. PubMed PMID: 29530936.

14. Lee YH, Martin-Orozco N, Zheng P, Li J, Zhang P, Tan H, Park HJ, Jeong M, Chang SH, Kim BS, Xiong W, Zang W, Guo L, Liu Y, Dong ZJ, Overwijk WW, Hwu P, Yi Q, Kwak L, Yang Z, Mak TW, Li W, Radvanyi LG, Ni L, Liu D, Dong C. Inhibition of the B7-H3 immune checkpoint limits tumor growth by enhancing cytotoxic lymphocyte function. Cell Res. 2017;27(8):1034–45. Epub 20170707. doi: 10.1038/cr.2017.90. PubMed PMID: 28685773; PMCID: PMC5539354.

15. Sun J, Fu F, Gu W, Yan R, Zhang G, Shen Z, Zhou Y, Wang H, Shen B, Zhang X. Origination of new immunological functions in the costimulatory molecule B7-H3: the role of exon duplication in evolution of the immune system. PLoS One. 2011;6(9):e24751. Epub 20110913. doi: 10.1371/journal.pone.0024751. PubMed PMID: 21931843; PMCID: PMC3172298.

16. Ling V, Wu PW, Spaulding V, Kieleczawa J, Luxenberg D, Carreno BM, Collins M. Duplication of primate and rodent B7-H3 immunoglobulin V- and C-like domains: divergent history of functional redundancy and exon loss. Genomics. 2003;82(3):365–77. doi: 10.1016/s0888-7543(03)00126-5. PubMed PMID: 12906861.

17. Fukushima A, Sumi T, Fukuda K, Kumagai N, Nishida T, Yamazaki T, Akiba H, Okumura K, Yagita H, Ueno H. B7-H3 regulates the development of experimental allergic conjunctivitis in mice. Immunol Lett. 2007;113(1):52–7. Epub 20070823. doi: 10.1016/j.imlet.2007.07.011. PubMed PMID: 17825429.

18. Nagashima O, Harada N, Usui Y, Yamazaki T, Yagita H, Okumura K, Takahashi K, Akiba H. B7-H3 contributes to the development of pathogenic Th2 cells in a murine model of asthma. J Immunol. 2008;181(6):4062–71. doi: 10.4049/jimmunol.181.6.4062. PubMed PMID: 18768862.

19. Yamato I, Sho M, Nomi T, Akahori T, Shimada K, Hotta K, Kanehiro H, Konishi N, Yagita H, Nakajima Y. Clinical importance of B7-H3 expression in human pancreatic cancer. Br J Cancer. 2009;101(10):1709–16. Epub 20091020. doi: 10.1038/sj.bjc.6605375. PubMed PMID: 19844235; PMCID: PMC2778545.

20. Mao L, Fan TF, Wu L, Yu GT, Deng WW, Chen L, Bu LL, Ma SR, Liu B, Bian Y, Kulkarni AB, Zhang WF, Sun ZJ. Selective blockade of B7-H3 enhances antitumour immune activity by reducing immature myeloid cells in head and neck squamous cell carcinoma. J Cell Mol Med. 2017;21(9):2199–210. Epub 20170411. doi: 10.1111/jcmm.13143. PubMed PMID: 28401653; PMCID: PMC5571514.

21. Cai D, Li J, Liu D, Hong S, Qiao Q, Sun Q, Li P, Lyu N, Sun T, Xie S, Guo L, Ni L, Jin L, Dong C. Tumor-expressed B7-H3 mediates the inhibition of antitumor T-cell functions in ovarian cancer insensitive to PD-1 blockade therapy. Cell Mol Immunol. 2020;17(3):227–36. Epub 20191014. doi: 10.1038/s41423-019-0305-2. PubMed PMID: 31611650; PMCID: PMC7051965.

22. Masemann D, Meissner R, Schied T, Lichty BD, Rapp UR, Wixler V, Ludwig S. Synergistic anti-tumor efficacy of oncolytic influenza viruses and B7-H3 immune-checkpoint inhibitors against IC-resistant lung cancers. Oncoimmunology. 2021;10(1):1885778. Epub 20210217. doi: 10.1080/2162402X.2021.1885778. PubMed PMID: 33643696; PMCID: PMC7894418.

23. Zhou Z, Yu X, Chen Y, Tan X, Liu W, Hua W, Chen L, Zhang W. Inhibition of the B7-H3 immune checkpoint limits hepatocellular carcinoma progression by enhancing T lymphocyte-mediated immune cytotoxicity in vitro and in vivo. Clin Transl Oncol. 2023;25(4):1067–79. Epub 20221213. doi: 10.1007/s12094-022-03013-4. PubMed PMID: 36512305.

24. Liu T, Gonzalez De Los Santos F, Rinke AE, Fang C, Flaherty KR, Phan SH. B7H3-dependent myeloid-derived suppressor cell recruitment and activation in pulmonary fibrosis. Front Immunol. 2022;13:901349. Epub 20220815. doi: 10.3389/fimmu.2022.901349. PubMed PMID: 36045668; PMCID: PMC9420866.

25. Du H, Hirabayashi K, Ahn S, Kren NP, Montgomery SA, Wang X, Tiruthani K, Mirlekar B, Michaud D, Greene K, Herrera SG, Xu Y, Sun C, Chen Y, Ma X, Ferrone CR, Pylayeva-Gupta Y, Yeh JJ, Liu R, Savoldo B, Ferrone S, Dotti G. Antitumor Responses in the Absence of Toxicity in Solid Tumors by Targeting B7-H3 via Chimeric Antigen Receptor T Cells. Cancer Cell. 2019;35(2):221–37 e8. doi: 10.1016/j.ccell.2019.01.002. PubMed PMID: 30753824; PMCID: PMC6645919.

26. Chapoval AI, Ni J, Lau JS, Wilcox RA, Flies DB, Liu D, Dong H, Sica GL, Zhu G, Tamada K, Chen L. B7-H3: a costimulatory molecule for T cell activation and IFN-gamma production. Nat Immunol. 2001;2(3):269–74. doi: 10.1038/85339. PubMed PMID: 11224528.

27. Mellacheruvu D, Wright Z, Couzens AL, Lambert JP, St-Denis NA, Li T, Miteva YV, Hauri S, Sardiu ME, Low TY, Halim VA, Bagshaw RD, Hubner NC, Al-Hakim A, Bouchard A, Faubert D, Fermin D, Dunham WH, Goudreault M, Lin ZY, Badillo BG, Pawson T, Durocher D, Coulombe B, Aebersold R, Superti-Furga G, Colinge J, Heck AJ, Choi H, Gstaiger M, Mohammed S, Cristea IM, Bennett KL, Washburn MP, Raught B, Ewing RM, Gingras AC, Nesvizhskii AI. The CRAPome: a contaminant repository for affinity purification-mass spectrometry data. Nat Methods. 2013;10(8):730–6. Epub 20130707. doi: 10.1038/nmeth.2557. PubMed PMID: 23921808; PMCID: PMC3773500.

28. Morley SC. The actin-bundling protein L-plastin: a critical regulator of immune cell function. Int J Cell Biol. 2012;2012:935173. Epub 20111213. doi: 10.1155/2012/935173. PubMed PMID: 22194750; PMCID: PMC3238366.

29. Todd EM, Deady LE, Morley SC. The actin-bundling protein L-plastin is essential for marginal zone B cell development. J Immunol. 2011;187(6):3015–25. Epub 20110810. doi: 10.4049/jimmunol.1101033. PubMed PMID: 21832165; PMCID: PMC3169714.

30. Todd EM, Deady LE, Morley SC. Intrinsic T- and B-cell defects impair T-cell-dependent antibody responses in mice lacking the actin-bundling protein L-plastin. Eur J Immunol. 2013;43(7):1735–44. Epub 20130518. doi: 10.1002/eji.201242780. PubMed PMID: 23589339; PMCID: PMC3794664.

31. Meadors JL, Cui Y, Chen QR, Song YK, Khan J, Merlino G, Tsokos M, Orentas RJ, Mackall CL. Murine rhabdomyosarcoma is immunogenic and responsive to T-cell-based immunotherapy. Pediatr Blood Cancer. 2011;57(6):921–9. Epub 20110401. doi: 10.1002/pbc.23048. PubMed PMID: 21462302; PMCID: PMC7401311.

32. Bruhns P. Properties of mouse and human IgG receptors and their contribution to disease models. Blood. 2012;119(24):5640–9. Epub 20120425. doi: 10.1182/blood-2012-01-380121. PubMed PMID: 22535666.

33. Wang Y, Kremer V, Iannascoli B, Goff OR, Mancardi DA, Ramke L, de Chaisemartin L, Bruhns P, Jonsson F. Specificity of mouse and human Fcgamma receptors and their polymorphic variants for IgG subclasses of different species. Eur J Immunol. 2022;52(5):753–9. Epub 20220225. doi: 10.1002/eji.202149766. PubMed PMID: 35133670.

34. Starbeck-Miller GR, Badovinac VP, Barber DL, Harty JT. Cutting edge: Expression of FcgammaRIIB tempers memory CD8 T cell function in vivo. J Immunol. 2014;192(1):35–9. Epub 20131127. doi: 10.4049/jimmunol.1302232. PubMed PMID: 24285839; PMCID: PMC3874719.

35. Baker M. Reproducibility crisis: Blame it on the antibodies. Nature. 2015;521(7552):274–6. doi: 10.1038/521274a. PubMed PMID: 25993940.

36. Bradbury A, Pluckthun A. Reproducibility: Standardize antibodies used in research. Nature. 2015;518(7537):27–9. doi: 10.1038/518027a. PubMed PMID: 25652980.

37. Roy AL, Wilder EL, Anderson JM. Validation of antibodies: Lessons learned from the Common Fund Protein Capture Reagents Program. Sci Adv. 2021;7(46):eabl7148. Epub 20211110. doi: 10.1126/sciadv.abl7148. PubMed PMID: 34757791; PMCID: PMC8580312.

38. Uhlen M, Bandrowski A, Carr S, Edwards A, Ellenberg J, Lundberg E, Rimm DL, Rodriguez H, Hiltke T, Snyder M, Yamamoto T. A proposal for validation of antibodies. Nat Methods. 2016;13(10):823–7. Epub 20160905. doi: 10.1038/nmeth.3995. PubMed PMID: 27595404; PMCID: PMC10335836.

